# MicroRNA function transitions from regulating developmental genes to transposable elements during the maturation of pollen

**DOI:** 10.1101/864017

**Authors:** Cecilia Oliver, Maria Luz Annacondia, Zhenxing Wang, R Keith Slotkin, Claudia Köhler, German Martinez

## Abstract

microRNAs play important roles to control the development of eukaryotic organisms. Both animal and plant microRNAs are essential for the spatio-temporal regulation of development but together with this role, plant microRNAs also control transposable elements and stimulate the production of epigenetically-active small interfering RNAs. This last role is evident in the plant male gamete containing structure, the male gametophyte or pollen grain, but how the dual role of plant microRNAs is integrated during its development is unknown. Here, we provide a detailed analysis of microRNA dynamics during pollen development and their genic and transposable element targets using small RNA and mRNA cleavage (PARE) high-throughput sequencing. Furthermore we uncover the microRNAs loaded in the two main Argonaute proteins in the mature pollen grain, AGO1 and AGO5. Our results indicate that the developmental progression from microspore to mature pollen grain is characterized by a reprogramming from microRNAs focused on the control of development to microRNAs regulating transposable element control.

## Main

Small non-coding RNAs control essential gene regulatory networks in eukaryotes at the transcriptional and postranscritional level. This broad term includes different classes of small RNAs (sRNAs) that have different biogenesis pathways, roles and cellular distribution but (in general) use sequence complementarity to recognize their target RNAs and silence their transcription and/or inhibit their translation^1^. The improvement of sequencing technologies has enabled to uncover the role of novel classes of sRNAs but also to understand better their cellular distribution and their roles during different stages of development of an organism or a tissue. According to their origin sRNAs can be classified in microRNAs (miRNAs), small interfering RNAs (siRNAs), piwi-interacting RNAs (piRNAs) or tRNA-derived sRNAs (tRFs), among others (for a recent review see:^2^).

In the case of plants, the sRNome is monopolized by two classes of sRNAs: siRNAs and miRNAs^3^. These two types of sRNAs have different biogenesis pathways and functions. siRNAs are the result of the processing of an RNA-DEPENDENT RNA POLYMERASE (RDR)-produced double stranded RNA by Dicer-like proteins (DCL), mainly DCL4, DCL2 and DCL3. This leads to the production of double stranded sRNAs of between 21 and 24 nucleotides (nts) of which one of the strands will be selectively incorporated into an Argonaute (AGO) protein^4^. On the other hand, miRNAs originate from *MIRNA* genes that produce non-coding transcripts with high self-complementarity that fold into a short hairpin structure. This hairpin is cleaved by DCL1 into a double-stranded sRNA of 21-22 nts in length. One of these sRNAs will then be selectively loaded into AGO1 and form the RISC complex, which uses the sRNA sequence to target mRNAs with perfect or imperfect sequence homology. In plants, this targeting normally leads to the cleavage of the mRNA, but can also induce translational repression^4,5^. Both miRNAs and siRNA regulate a diversity of processes including development, defense, reproduction and genome stability. However, generally, siRNAs regulate heterochromatin and development/defense via DNA methylation and secondary sRNAs respectively, while miRNAs are associated with the regulation of development through the posttranscriptional control of transcription factor mRNAs ^4^. Nevertheless these two sRNA classes are intertwined in certain aspects of development. For example, regulation of auxin signaling by the generation of trans-acting siRNAs is coordinated by miRNAs^6^. In plants, miRNAs also play a role in genome protection through the initial targeting of transposable elements (TEs) and generation of secondary siRNAs from their transcripts^7,8^. Interestingly, one of the miRNAs targeting TEs, miR845, is involved in the generation of TE-derived siRNAs in the male gametophyte and mediate genome dosage response^9^. This diversity of miRNA-regulated processes highlights their elasticity as regulatory molecules.

The functional versatility of miRNAs is especially important during reproduction, where cells face the duality of carrying out a very specific developmental program. In other organisms like zebrafish^10^, mouse^11^, *C. elegans*^12^ or *D. melanogaster*^13^ miRNAs do not only have a differential accumulation pattern in sperm, but play important roles in sperm maturation, fertilization and post-fertilization events. However, little is known about how miRNA activity might shape the transcriptome during pollen development. Strong *ago1* and *dcl1* mutants have different reproductive abnormalities and reduced seed set^14-16^, which points to an important role of miRNAs during reproduction in plants.

Previous reports of *Arabidopsis* pollen sRNAs have focused on the analysis of the accumulation of these only in the mature pollen grain^16,17^. Here we analyze in depth the contribution of miRNAs to the sRNA population during the different stages of pollen grain development, their loading into AGO proteins and their target mRNAs. Overall, our work suggests that miRNAs involved in epigenetic regulation, like miR845, are enriched at later stages of pollen grain development correlating with their preferential loading in AGO5. In contrast, miRNAs targeting mRNAs from genes involved in development decrease their accumulation during pollen development. This coincides with increased expression of their target genes at pollen maturity, which are mainly involved in pollen grain germination. Additionally, we identify a group of miRNA-regulated TEs in the pollen grain. In summary, this work shows that miRNAs modulate both the transcriptional and epigenetic reprogramming of the pollen grain.

## Results

### Dynamic accumulation of miRNAs during pollen development

To understand the potential changes in development during the transition leading to the mature pollen grain, we focused on analyzing miRNA accumulation at four different stages of pollen development (uninuclear, binuclear, trinuclear and mature pollen grain, representative pictures shown in Figure 1a and Supplementary Figure 1). Using density centrifugation we isolated four different developmental stages of pollen as previously described ^18^. We isolated total RNA, prepared and sequenced sRNA libraries from two biological replicates for each of these stages (Supplementary Figure 1 and Supplementary Table 1). The total miRNA accumulation profiles between the different developmental stages did not reveal striking differences between the stages (Figure 1b); we only found a slight increase in 22 nt miRNAs in trinuclear and mature pollen grain in comparison to uni- and binuclear (Figure 1b). The analysis of qualitative differences in the miRNA populations between our samples (Figure 1c) revealed that the majority of miRNA families are present in all our libraries (233 miRNAs); however, we also detected specific miRNAs in each developmental stage, including 23 miRNAs in the mature pollen grain and 13, 7, and 5 in the uni-, bi- and trinuclear stages, respectively (Supplementary Table 2). We further analyzed the quantitative changes experienced by the most highly accumulating miRNAs (Figure 1d). The comparison between the accumulation level of the top seven accumulating miRNA families (representing more than 90% of all sRNAs in uni-, bi- and tricellular pollen and 77% of the mature sRNAs) revealed that there is a striking progressive decrease in the accumulation of some miRNA families during pollen development. In particular, miRNAs involved in developmental processes, like miR156, miR161.1, miR159, miR166, and miR162 (Figure 1d) decrease at later stages of pollen development. In comparison, the relative levels of miR845 increase substantially during pollen development until occupying close to 45% of the all sRNAs at pollen maturity (Figure 1d). We confirmed this decrease in the accumulation of miRNAs for miR156 and miR161.1 by Northern blot (Figure 1e). Together, our analysis shows that during pollen development there is a transition from a diverse miRNA pool to a miRNA pool monopolized by miR845.

**Figure 1.**
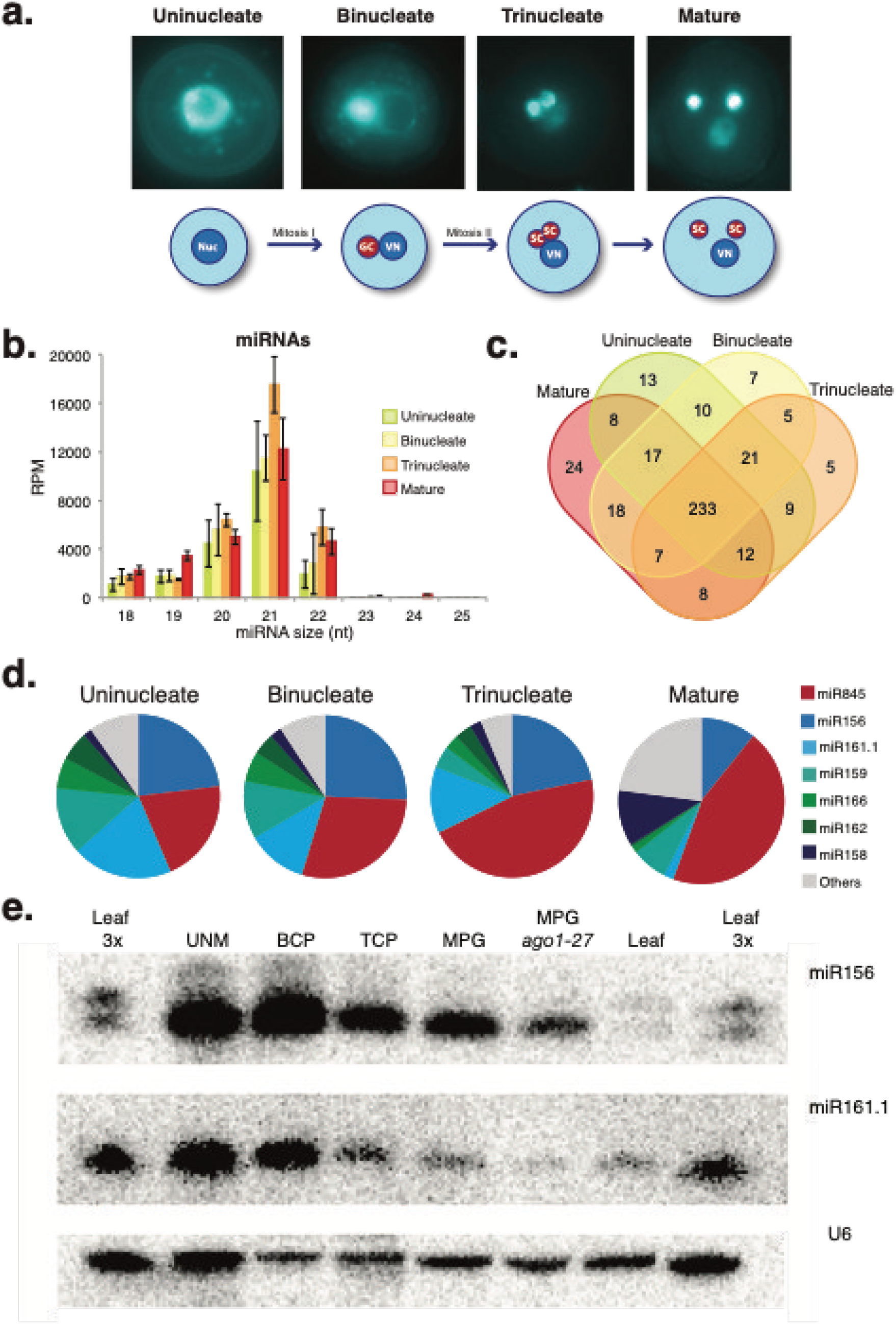
miRNome dynamics during pollen development: a) Representative images of the pollen developmental stages analyzed here. b) microRNA size distribution and accumulation for the developmental stages shown in a. c) Venn diagram showing the common and developmental stage-specific miRNAs for the stages indicated. d) Pie charts depicting the accumulation of main miRNAs during pollen development for each developmental stage. e) Northern blot showing the decrease in accumulation of two developmentally related miRNAs miR156 and miR161.1. The U6 small nuclear RNA was used as a control for RNA loading.

### AGO1 and AGO5 loading explains miRNA enrichment in the mature pollen grain

miRNA loading into AGO proteins determines their effect at the cellular level ^19^. Several AGO proteins have been reported to be active in the male gametophyte, including AGO1, AGO2, AGO4, AGO5, and AGO9 ^16,20-22^, but the sRNA populations loaded into them have not been studied. To shed light into the characteristics of the RISC complexes in the pollen grain, we analyzed the sRNAs bound to the main miRNA-related AGO proteins expressed in the pollen grain: AGO1 and AGO5 (Supplementary Figure 2). In the mature pollen grain, these two AGOs have a different cellular localization; while AGO1 is located in the vegetative nucleus and in the vegetative cell (VC)^23^, AGO5 accumulates in the sperm cell (SC) cytoplasm^16^. We investigated the cellular localization of both AGOs during pollen grain development and found that AGO1 is present already in the cytoplasm of unicellular pollen at the uninuclear stage and this expression pattern is maintained in the VC until the mature stage (Figure 2a). On the other hand, AGO5 was only detectable in the GC/SCs at the late binuclear/early trinuclear stage (Figure 2a). Next, we identified the sRNAs loaded into both AGOs by sequencing of sRNAs that were immunoprecipitated using AGO-specific antibodies. In line with their predicted role, we detected enrichment for miRNAs in the immunoprecipitated sRNAs compared to their input controls (Figure 2b). Both AGOs shared a proportion of their respective miRNomes (54.5% and 61.3% for AGO1 and AGO5, respectively, Figure 2c), in particular both had a strong preference to load miR845 family members (37.3% and 71.3% of the total miRNome for AGO1 and AGO5 respectively). AGO1 also loaded a substantial fraction of miRNAs with well-known roles in developmental processes, like miR158 (12.4%), miR159 (9.5%), miR156 (8.5%), miR403 (6%), and miR168 (4.8%), while AGO5 loaded only a small fraction of developmental-related miRNAs, like miR156 (5.9%) or miR158 (5.6%) (Figure 2d). This different miRNA-loading pattern might reflect the different roles of both AGOs in relation to their cellular localization, with AGO1 being required to regulate the developmental program and post-transcriptional activity of TEs in the VC ^20,24^, AGO5 might mediate specifically TE control in the SCs ^9^. This correlates with the known role of the VC in the regulation of pollen development and germination^25^. In summary, both, the cellular localization of AGO1 and AGO5 and their loaded miRNAs correlate with two different programs mediating the regulation of development and TEs respectively.

**Figure 2:**
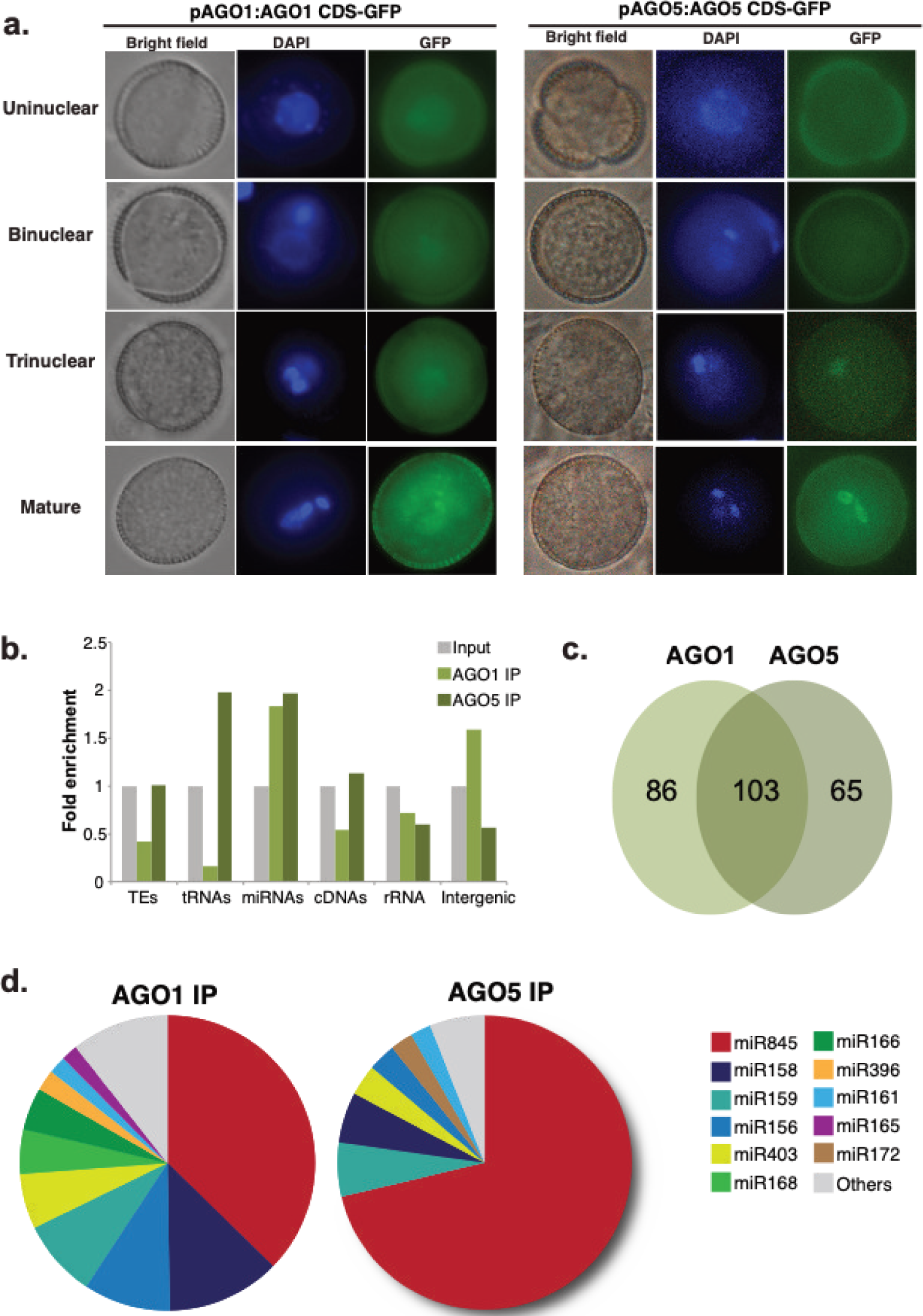
Comparison of AGO1 and AGO5 loaded miRNAs. a) Analysis of the cellular localization of AGO1 and AGO5-GFP fusion proteins during pollen development. b) Analysis of the enriched categories for sRNAs between 18 and 28 nts for AGO1 and AGO5 immunoprecipitated sRNAs compared to their respective input control. c) Venn diagram showing the number of common and exclusive miRNAs for each AGO protein. d) Pie chart depicting the miRNAs loaded preferentially on each of the AGO proteins under study.

### Inhibition of miRNA activity affects pollen development

To evaluate the level of influence of miRNAs on pollen grain development, we aimed to inhibit their activity at late stages of pollen development where AGO1 and AGO5 primarily accumulate (Figure 2a). Strong AGO1 mutant alleles fail to develop viable gametes^14^. We hypothesize that overexpression of a viral silencing suppressor using specific promoters would drive a cell-specific reduction in AGO1 activity as previously reported^20^. The Tombusvirus silencing suppressor P19 is a well-studied protein that inhibits miRNA/miRNA* duplex action^26^. Accordingly, we expressed P19^27^ fused to RFP under the control of the *KRP6* promoter to drive the expression at late stages of development of the pollen grain VC^28^ (Figure 3a). *KRP6pro::P19-RFP* transgenic lines had defects in pollen development with many of the mature pollen grains aborted at maturity (Figure 3b) and inhibition of pollen grain germination (Figure 3c). In brief, inhibition of miRNA activity in the male gametophyte VC supports a role of VC miRNAs in the posttranscriptional regulation of genes required for pollen development.

**Figure 3:**
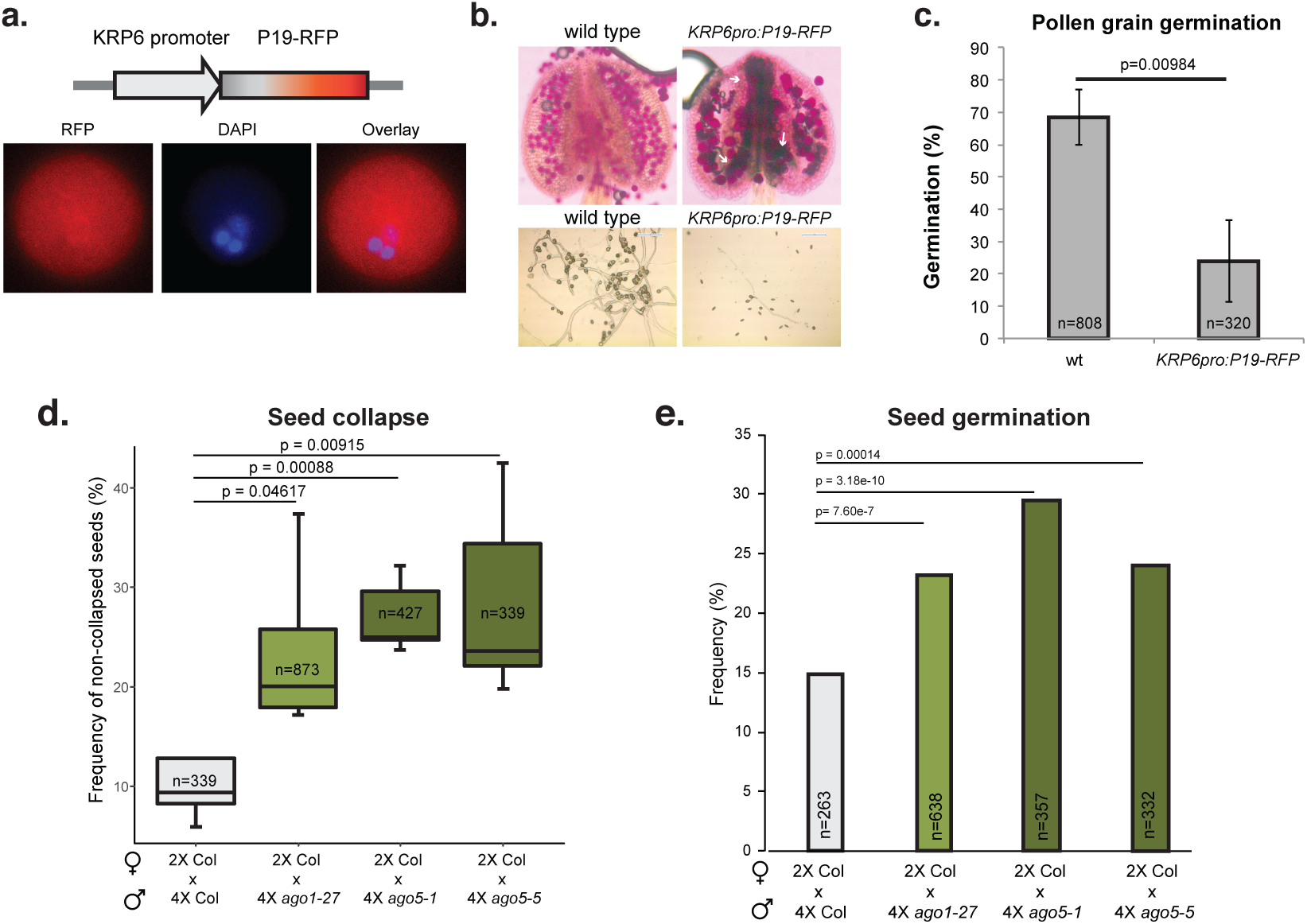
Inhibition of miRNA activity in the pollen grain leads to developmental defects of pollen grain development and inhibition of germination. a) Diagram of the construct used to express the viral silencing suppressor P19 in the mature pollen grain and representative image of its accumulation in mature pollen grains. b) Representative pictures of the analysis of pollen defects by Alexander staining and in vitro germination for wt and P19 transgenic plants. Aborted pollen grains clutches are indicated with white arrows. c) Pollen grain germination percentages for wt and KRP6pro:P19-GFP transgenic line. Number of individual pollen grain measurements (n) is shown inside of each bar. Error bars represent the standard deviation values for the three bioreplicates analyzed. P value is the result of a standard t-test with 2 tails and unequal variance. d) Frequency of non-collapsed and e) germinated seeds derived from crosses of wt (2xCol) maternal parents with 4x mutants of indicated genotypes. T-test and Chi-squared test were performed in E and F, respectively. Number of individual seed measurements (n) is shown inside of each bar. Whiskers in the box plots extent to the maximum and minimum values.

### AGO1 and AGO5 are required for the triploid block response

Gametic sRNAs establish hybridization barriers in different species ^29,30^. In plants, polyploidization establishes hybridization barriers due to unbalanced expression of imprinted genes in the endosperm ^31-35^, in a phenomenon known as the triploid block^36^. In *Arabidopsis*, the triploid block is established upon crosses of a pollen donor forming 2n pollen with a diploid maternal plant. Depletion of the major Pol IV subunit *NRPD1A* or the miRNA gene *MIR845B* suppresses the triploid block response ^9,37-39^. To test whether the miRNA populations loaded in AGO1 or AGO5 are responsible for establishing the triploid block, we created tetraploid versions of the AGO1 weak allele *ago1-27* and of the AGO5 alleles *ago5-1* and *ago5-5* and performed crosses with a wt diploid mother. The results of those pollinations revealed that paternal tetraploid *ago1-27* weakly, but nevertheless significantly increased triploid seed viability (Figure 3d and e). Similarly, paternal tetraploid *ago5-1* and *ago5-5* significantly increased triploid seed viability and seed germination (Figure 3d and e) to a similar level as *ago1*, suggesting that both redundantly function in the triploid block response. Thus, paternal AGO1 and AGO5 are part of the triploid block response potentially through their loading of microRNA family members.

### Genome-wide analysis of miRNA-regulated transcripts in the pollen grain

To uncover miRNA-regulated transcripts in the pollen grain, we generated and sequenced PARE libraries from mature pollen grains from Col-0 wild type plants and from mature rosette leaves, as previously described ^40^ (Supplementary Table 1). PARE is a technique that targets cleaved mRNAs with a polyA tail but without a 5’ cap for library preparation and sequencing. A comparison of miRNA-targeted mRNAs between leaf and pollen highlighted the tissue specificity of miRNA regulation, as only 21.5% of pollen miRNA-targeted genes were also miRNA-regulated in the leaf (Fig 4a). These target mRNAs are regulated by miRNAs that are both shared with leaf (68% of all pollen miRNAs) and pollen-specific (representing 32% of the total active miRNAs). A global analysis of pollen miRNA-regulated transcripts revealed that the majority are associated with developmental processes related to transport, cell organization and biogenesis, signal transduction, and response to stress (Supplementary Figure 4 and Supplementary Table 3).

**Figure 4:**
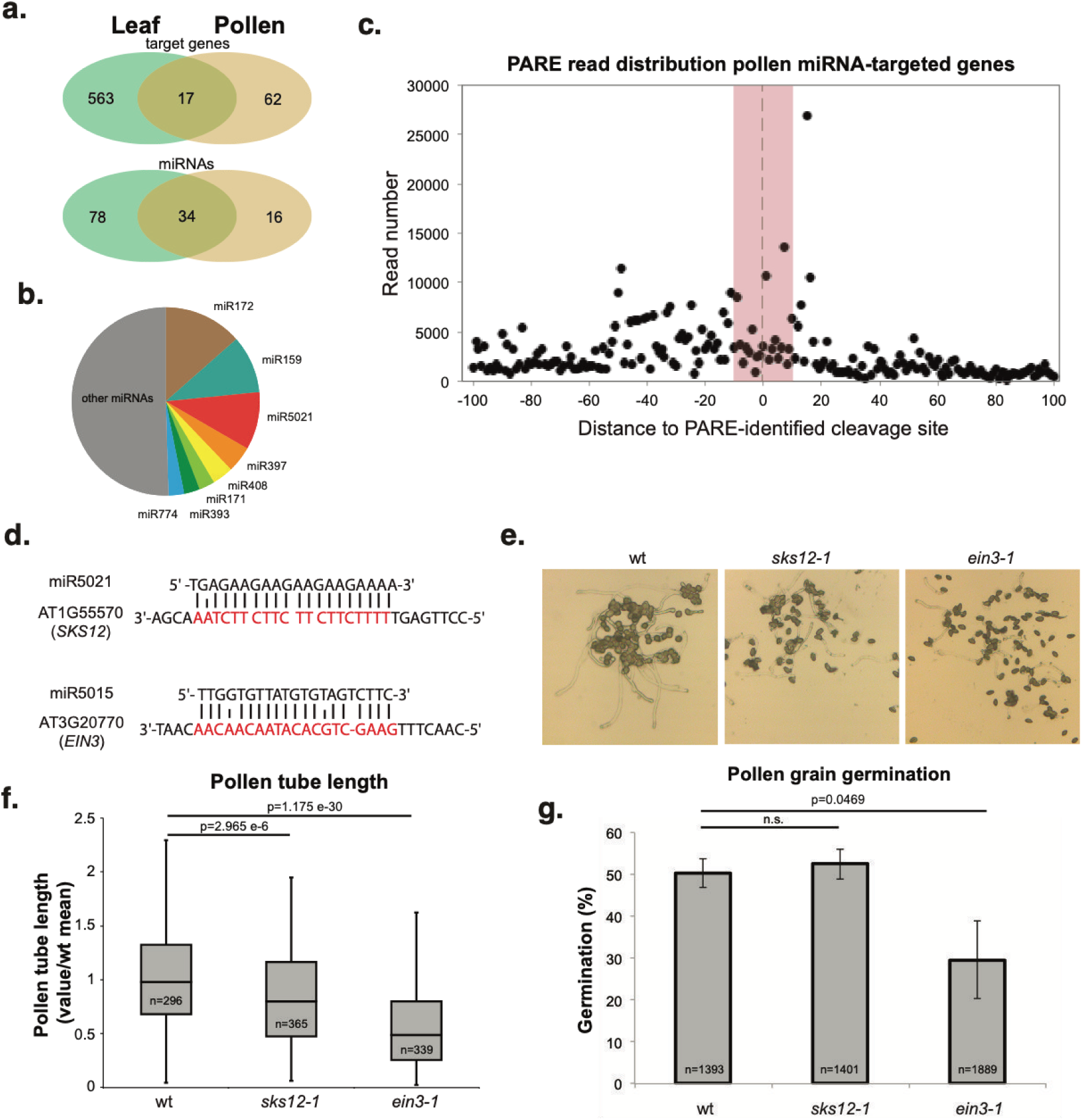
Analysis of miRNA genic targets in the mature pollen grain identified by PARE sequencing: a) Venn diagrams showing the common and tissue specific target genes and active miRNAs for the tissues analyzed. b) Pie chart showing highly represented miRNA target sites on PARE confirmed miRNA target genes. c) Distribution of 5’ ends of PARE reads around the predicted cleavage site (located at coordinate 0 in the X axis) in a 100 nt window. Red zone represents the physical position covered by the bound miRNA. d) Examples of two miRNA targets in our PARE analysis: SKS12-miR5021 and EIN3-miR5015. e) Representative pictures of pollen grain germination for wt and the *sks12-1* and *ein3-1* mutants. f) Length of the pollen tube and g) percentage of germination for in vitro germinated pollen grains for the genotypes indicated. Number of individual pollen grain measurements (n) is shown inside of each bar. Error bars in h represent the standard deviation values for the three bioreplicates analyzed. P value is the result of a standard t-test with 2 tails and unequal variance. Whiskers in the box plots extent to the maximum and minimum values.

Pollen miRNA target mRNAs are involved both in pollen grain development and pollen tube germination and include well-known regulators of these processes such as *SK32, AtbZIP34* and *AFB3* involved in pollen development^41-43^, or *MYB97, MYB101*, and *SYP131* involved in pollen tube germination^44,45^. miRNAs with a higher number of targeted transcripts included juvenile-to-adult phase transition related miR172^46^ and miR159^47^ (13.5 and 10% of targeted genes) and also the pollen specific miR5021 (10% of targeted genes) (Figure 4b). miRNA targets included classic miRNA-regulated genes, such as the *TAS* genes or miR172-targeted transcription factors *APETALA2 (AP2)* and *TARGET OF EARLY ACTIVATION TAGGED (EAT) 2 (TOE2)* (Supplementary Figure 4). We also detected pollen-specific targeting events like the targeting of *SKU5 SIMILAR 12* (*SKS12)* by miR5021 or *ETHYLENE-INSENSITIVE3* (*EIN3*) by miR5015 (Supplementary Figure 3 and Supplementary Table 4). In other organisms, miRNA targeting in the gametes increases the stability of targeted transcripts ^48^. We explored if a similar scenario may apply to *Arabidopsis*. The distribution of 5’-P end reads in a 100 nt window from the predicted target site shows that most of these targeting events resulted in cleavage of the target RNAs without evidence of ribosomal stalling (Figure 4c). All these evidences show that miRNA activity in the pollen grain induces the cleavage of transcripts involved in pollen grain development and germination.

To test this observation, we obtained homozygous mutants for two of the genes specifically regulated by miRNAs in pollen identified in our PARE analysis: *SKS12* (AT1G55570, mutant termed *sks12-1*, Supplementary Figure 4) and *EIN3* (AT3G20770, *ein3-1)*^49^ (Figure 4d) and evaluated the ability of their mature pollen grain to germinate *in vitro* (Figure 4e). Measurement of pollen tube length and germination rate after 16 hrs of incubation indicated that while only *ein3-1* was affected in the rate of pollen germination (Figure 4g), both mutants were impaired in pollen tube growth (Figure 4f). Thus, we conclude that miRNAs regulate developmental processes that are important for pollen development and pollen grain germination.

### miR845-targeted TEs progressively decrease their level of 24 nt sRNAs during pollen development

miRNAs have been identified as important posttranscriptional regulators of TEs^7,9^. In particular, the miR845 family is involved in the biogenesis of epigenetically-active siRNAs (easiRNAs) through the targeting of the primer binding site (PBS) of TEs^9^. This miRNA family is composed by two members, miR845a and miR845b, which are 21 and 22 nts in length, respectively ^9^. Analysis of their presence in our pollen development sRNA libraries showed that, during pollen development, the two members of the miR845 family increased their accumulation, especially miR845a which increased by 2.7 fold (Figure 5a). Both miRNAs did not seem to be affected by fluctuations of the processing precision and were of the expected size at all stages of development (Supplementary Figure 5). Interestingly, the 5’ terminal nucleotide of miR845a and b (C and T respectively, Figure 5b) suggests a preferential loading in AGO5 and AGO1, respectively ^50^. We analyzed if this predicted differential loading was detectable in our AGO1 and AGO5 IP sRNA libraries and, indeed, AGO5 showed a clear preferential loading of miR845a (62% of AGO5 IPed sRNA sequences, Figure 5c).

**Figure 5:**
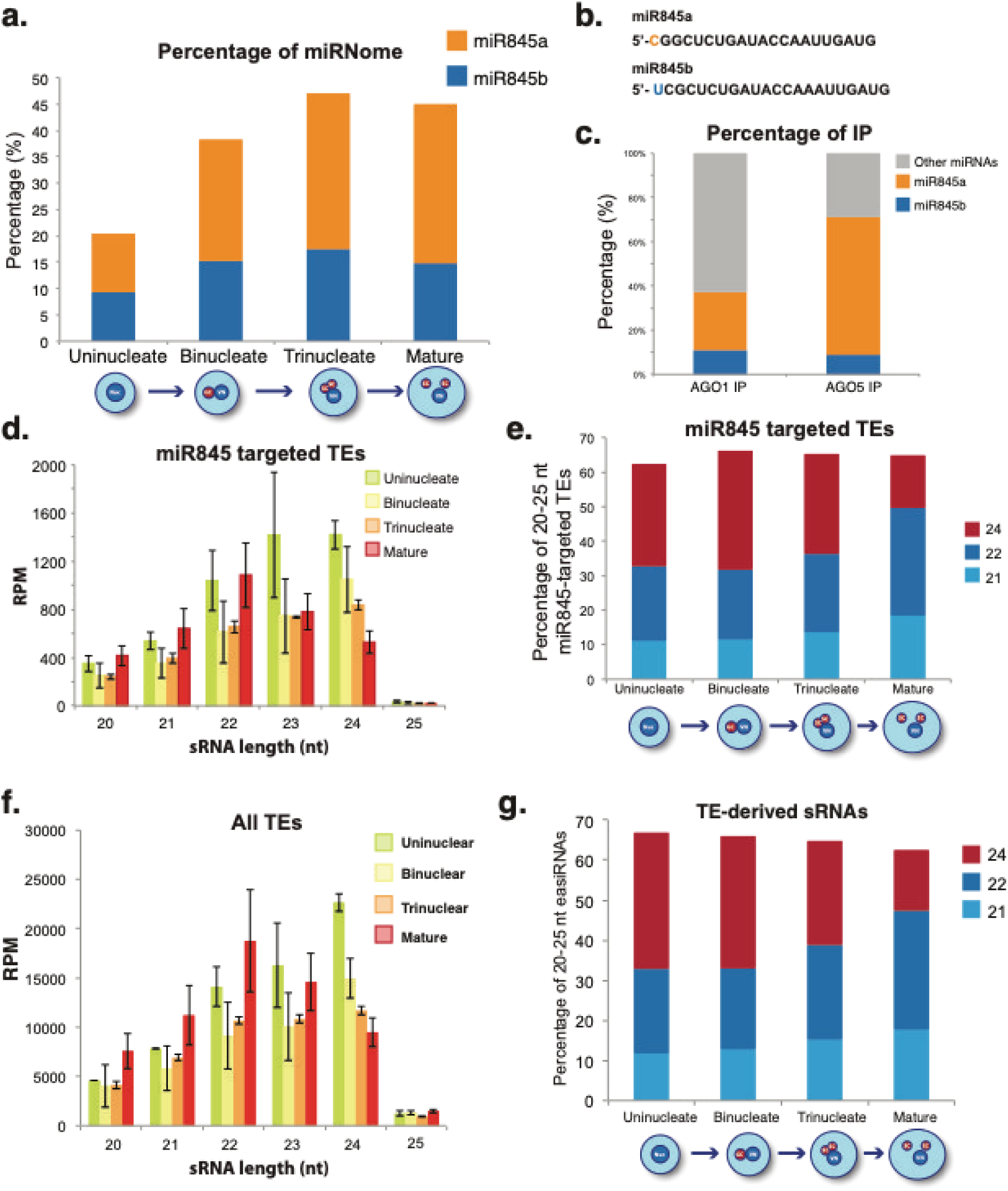
Analysis of miR845 dynamics and its target TEs during pollen development. a) Accumulation percentages for miR845a and b during pollen development. b) Sequence comparison of miR845a and b with the 5’nucleotide highlighted. c) Percentage of the impact of miR845 a and b on total AGO1 and AGO5 immunoprecipitated miRNAs. d) Accumulation size profile of TE-derived siRNAs of predicted miR845-targeted TEs. e) Accumulation of 21,22 and 24 nt sRNAs from miR845-targeted TEs during pollen development shown as percentages of total 20-25 nt sRNAs derived from those TEs. f) Accumulation profile of all TE-derived sRNAs. g) Accumulation of 21,22 and 24 nt sRNAs from TEs during pollen development shown as percentages of total 20-25 nt easiRNAs.

Members of the miR845 family were prosed to trigger Pol IV-dependent easiRNA biogenesis during meiosis or early gametogenesis^9^. Analysis of easiRNAs derived from miR845-targeted TEs indicates that easiRNAs accumulate to high levels already at the unicellular stage (Figure 5d). As expected from its preferential loading in AGO1, miR845b targeted-TEs produce easiRNAs earlier and to a greater extent than miR845a targets (Supplementary Figure 5). Interestingly, during pollen grain development there is a gradual transition from a majority of 24 nts at the unicellular stage to a majority of 22 nts at pollen maturity (Figure 5d-e). This shows that, most probably, miR845-dependent easiRNA biogenesis takes place after meiosis progressively during the two rounds of pollen mitosis. This tendency of losing 24 nt sRNAs during pollen grain development and gaining 21/22 nt sRNAs is common for all TEs (Figure 5f-g), revealing that similar mechanisms to miR845 targeting might exist for all TEs. Unfortunately, TE-targeted by miR845 could not be confirmed using our PARE sequencing (Supplementary Table 4) since PARE libraries are prepared from polyadenylated transcripts^40^ and miR845 targets non-polyadenylated Pol IV transcripts^9^. This fact is in agreement with the prediction that miR845 family members target exclusively Pol IV transcripts^9^. Overall, these results show that TEs tend to loss 24 nts while gaining 21/22 nt sRNAs during pollen development in correlation with an increase in the accumulation of miR845 family members.

### PARE sequencing analysis identified a pool of Pol II-transcribed and miRNA-regulated TEs

Although our PARE libraries could not be used to identify miR845-targeting events, they allowed the identification of miRNA-targeted polyadenylated TE transcripts. We identified a number of miRNA-targeted TEs (Supplementary Table 4), strongly suggesting that Pol II transcribes these TEs. Interestingly, these TEs were targeted mostly by pollen-specific miRNAs like miR5021, miR5658 and miR5645 (that together represent 58% of the targeting events identified) (Figure 6a). Similar to the targeting of genic mRNAs, distribution of reads in a 100 nt window from the PARE-identified cleavage site for TEs revealed a clear preference for RNA cleavage (Figure 6b). Most of these TEs localized to euchromatic regions (57.5%, Figure 6d) and belonged to the MuDR, Copia, En-Spm and Gypsy families (82% of total miRNA-targeted TEs, Figure 6c). miRNA-targeted TEs dramatically lost 24 nts during the transition from unicellular to bicellular pollen (Figure 6e), revealing that their regulation is different from the rest of TEs (Figure 5f).

**Figure 6:**
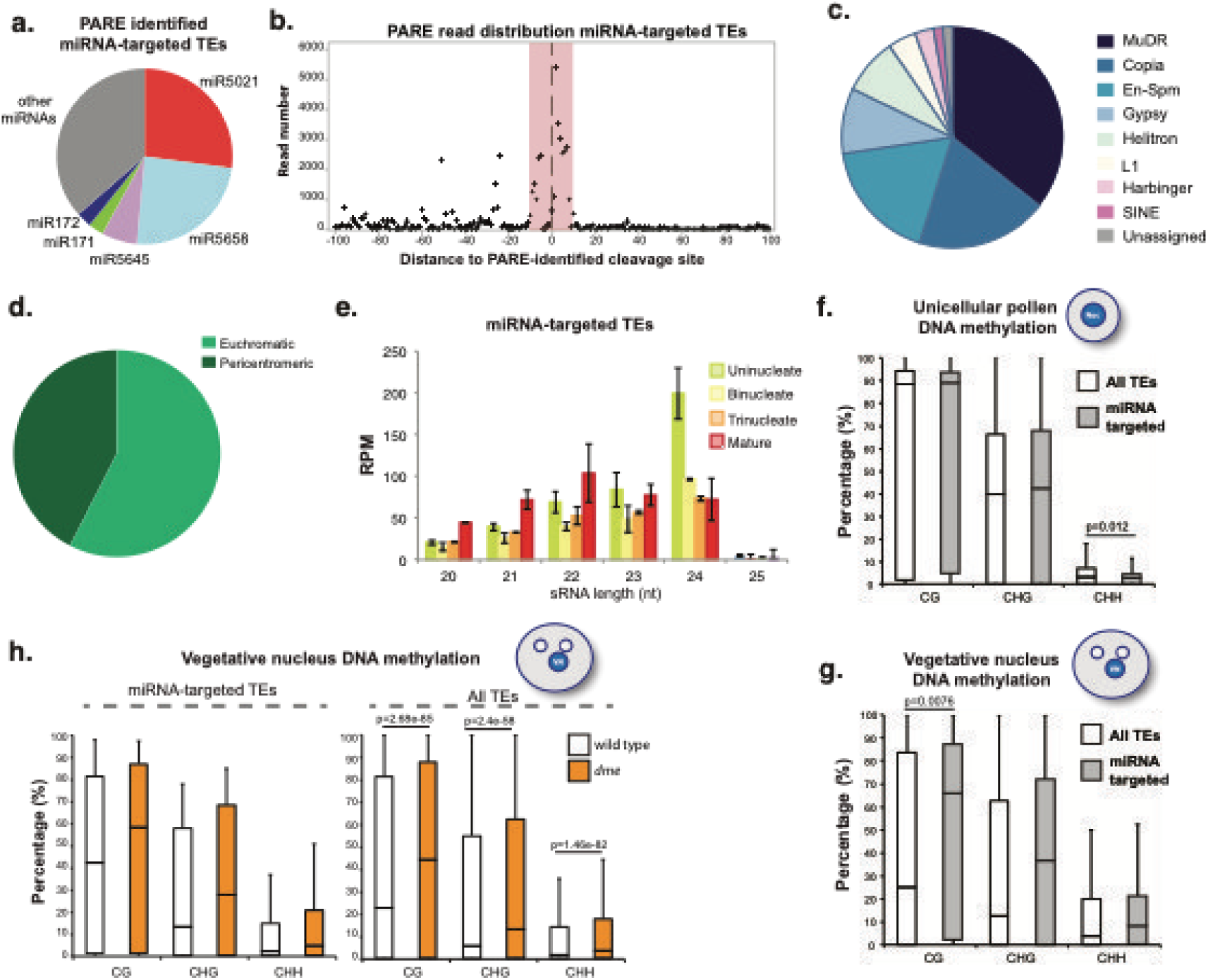
Analysis of miRNA TE targets in the mature pollen grain identified by PARE sequencing. a) Pie chart showing highly represented miRNA target sites on PARE confirmed miRNA-targeted TEs. b) Global distribution of 5’ ends of PARE reads around the predicted cleavage site for TEs (located at coordinate 0 in the X axis) in a 100 nt window. Red zone represents the physical position covered by the mRNA-bound miRNA. c) Family categorization and d) genomic distribution of miRNA-targeted TEs. e) sRNA accumulation size profile for TEs predicted to be targeted by miRNAs and transcribed by Pol II. f-g) Levels of cytosine methylation for the different contexts (CG, CHG and CHH) in the unicellular pollen grain (f) and vegetative nucleus (g) for all TEs (white boxes) or miRNA-targeted TEs (grey boxes). h) Levels of cytosine methylation for the different contexts (CG, CHG and CHH) in the vegetative nucleus for miRNA-targeted TEs (left panel) or all TEs (right panel) in wild type (white boxes) and *dme* (orange boxes). In all graphs, P value is the result of a standard t-test with 2 tails and unequal variance. Only significant differences between measurements are highlighted in the graphs. Whiskers in the box plots extent to the maximum and minimum values.

Next, we analyzed the levels of DNA methylation of PARE-identified miRNA-targeted TEs during pollen development^51^ and its potential connection to their sRNA levels using publicly available data (Supplementary Table 5). miRNA-targeted TEs have significantly lower levels of CHH methylation at the unicellular stage compared to the rest of TEs (Figure 6f), pointing to their dependence on the RdDM pathway to retain sRNA-based CHH methylation. At the mature developmental stage, these TEs retain significant higher levels of CG methylation in the VN compare to other TEs (Figure 6g) while maintain low levels of CHH methylation in the SCs (Supplementary Figure 6). Altogether this may indicate that this group of TEs is not a target of DME-mediated demethylation in the VN. Subsequently, analysis of DNA methylation in the VN of *dme* mutants confirmed that miRNA-targeted TEs are indeed not targeted by DNA glycosylases (Figure 6h), which translates into maintenance of low CHH methylation in the SCs (Supplementary Figure 6). Altogether, our data points to the existence of Pol II-transcribed TEs during pollen epigenetic reprogramming that are not targeted by DME but are regulated by miRNAs.

### The miRNome adjusts during pollen development to adapt the regulation of genic and TE targets

Finally, to understand the role of the identified active miRNAs during pollen development, we studied in detail their accumulation patterns in our sRNA libraries from different stages of pollen development. During pollen maturation, the majority of miRNAs regulating genes and TEs (and the common miRNAs between both groups, Supplementary Figure 7a) maintained or decreased their level of accumulation (65.7%, Figure 7a and Supplementary Figure 7b-c). This decrease was especially evident at the tricellular stage where the SCs appear in the VC and AGO5 accumulation was evident (Figure 2A). Interestingly, the members of the miR845 family followed the opposite trend, with a progressive accumulation during pollen development (Figure 7a). In parallel with the decrease of Pol II-active miRNA accumulation, their genic targets increased their expression towards maturity of the pollen grain (Figure 7c). The level of expression of these miRNA-target genes was even maintained during pollen grain germination (Figure 7c), indicating that miRNAs regulating genic products in the pollen grain have an important role in the overall regulation of transcripts available for pollen development and germination.

**Figure 7.**
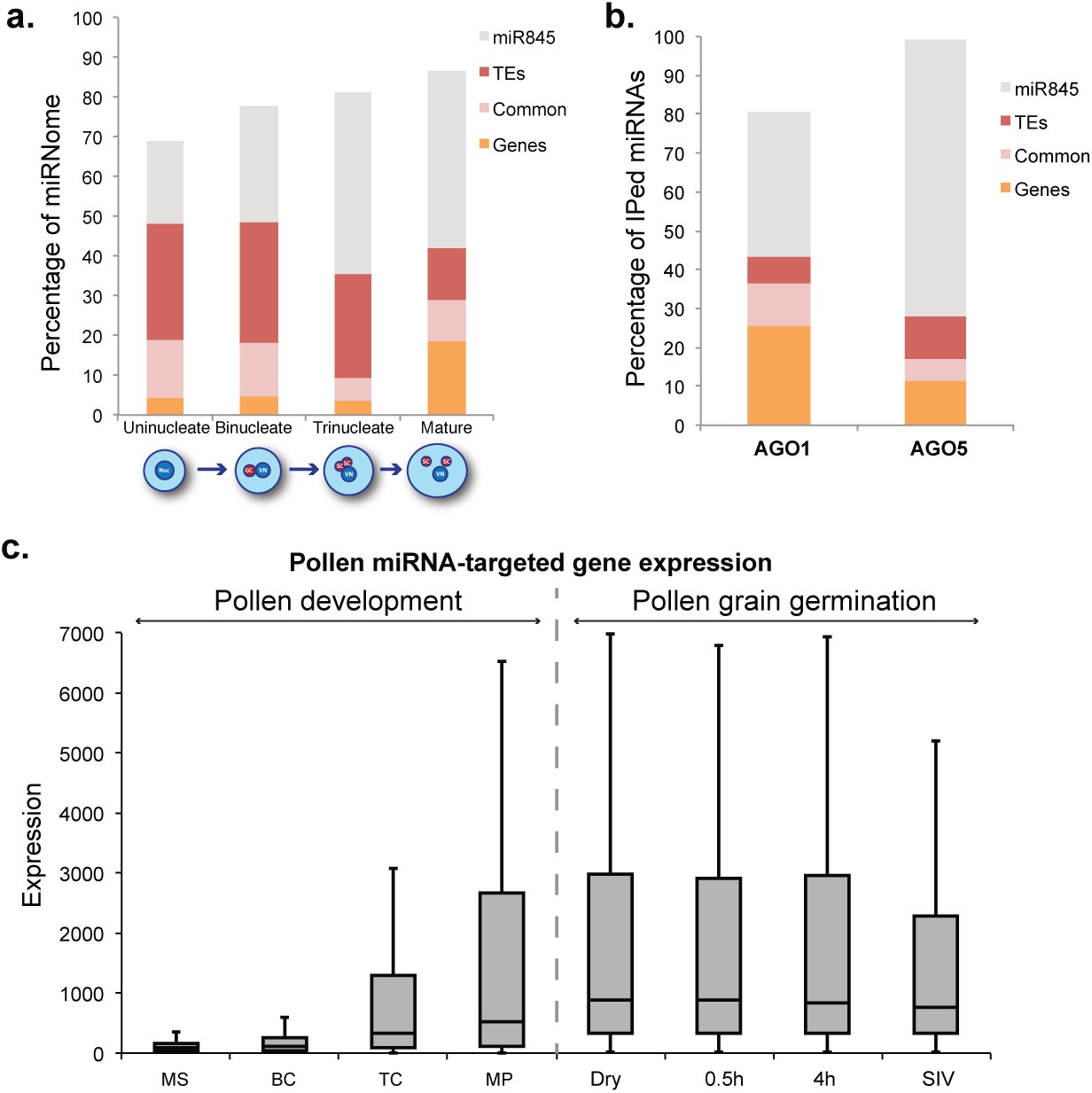
a) Distribution of miR845 family and PARE-identified miRNAs targeting TEs, miRNAs or both (termed common) during pollen development. Accumulation values are represented as percentage of the total miRNome. b) Presence of miR845 family and PARE-identified miRNAs targeting TEs, miRNAs or both (termed common) in AGO1 and 5 immunoprecipitated sRNAs. c) Level of expression of miRNA target genes during pollen development and pollen grain germination present in the ATH1 microarray (MS= microspore, BC=bicellular pollen, TC= tricellular pollen, MP= mature pollen grain, Dry= Dessicated mature pollen, 0.5h= In vitro-germinated pollen grains after 30 minutes, 4h= In vitro-germinated pollen grains after 4 hours and SIV= Pollen tubes grown through the stigma and style). Whiskers in the box plots extent to the maximum and minimum values.

The analysis of the two populations of active miRNAs present in our AGO immunoprecipitated libraries helped to understand to a better extent their role during pollen development (Figure 7b). AGO1 tended to load a mix of miRNAs involved both in the regulation of development (36.6%) and TEs (44 %) (Figure 7b). On the other hand AGO5 loads a majority of miRNAs involved exclusively in the regulation of TEs (82%, Figure 7b). In summary, overall, our analysis shows that during pollen development there is a transition from a diverse miRNA pool that regulates both development and Pol II-transcribed TEs, probably loaded in AGO1, to a miRNA pool that, at maturity, controls Pol IV-transcribed TEs monopolized by miR845 and with preferential loading in AGO5.

## Discussion

Through the use of pollen stage separation combined with high-throughput sRNA sequencing, PARE sequencing and the characterization of several characteristics of pollen grain development, we have (1) identified the characteristics of the miRNome during pollen grain development, (2) determined the miRNA populations loaded into the main AGO proteins in the pollen grain, AGO1 and AGO5, (3) identified miRNA targets (both TEs and genes) and (4) identified the involvement of both AGO1 and AGO5 in the triploid block. Our data reveal that the miRNome experiences a reprogramming during pollen development, transitioning from a miRNome mainly involved in developmental control to a miRNA population focused on the transcriptional control of TEs.

The pollen grain undergoes both a transcriptional and epigenetic reprogramming during its transition to maturity, but whether the first is a consequence of the latter is unknown ^24,51,52^. Our data indicates that miRNAs also experience a reprogramming, which could influence the transcriptional and epigenetic changes taking place in this tissue. Reduction of miRNAs in the pollen grain via the expression of the P19 viral silencing suppressor strongly reduced pollen grain viability and germination (Figure 3). Indeed our identification of miRNA target mRNAs in the pollen grain of *Arabidopsis* (Figure 4 and 6) through PARE sequencing shows their importance in the regulation of both genes and TEs.

In this tissue, genes targeted by miRNAs had a higher level of expression in the mature pollen stage and were involved in processes related with pollen germination (Figure 7c). Although this might suggests that gametic miRNAs increase the stability of transcripts, similar as previously observed in *C*.*elegans*^48^, our PARE data indicates that most probably the miRNA targeting events identified here induce the cleavage of their target mRNAs (Figure 4c), showing that developmental miRNA targets in pollen are expressed at high levels and miRNA targeting dampens their expression (Figure 7a-c, model shown in Supplementary Figure 8). As a proof of concept, we analyzed the defects in pollen grain germination experienced by two miRNA target genes identified in our analysis: *SKS12* and *EIN3* (Figure 4d-g).

Additionally we have explored the influence of the miRNome on the epigenetic regulation of TEs in the pollen grain. During pollen grain development the miR845 family members (miR845a and b) increase in abundance (Figure 1d). This increase likely results in the preferential loading of these two miRNAs in AGO5, a highly abundant AGO protein in the sperm cells (Figure 2a). miR845 members have been proposed to target Pol IV transcripts of several retrotransposons and induce the production of 21/22 easiRNAs from those transcripts^9,37^. Our analysis indicates that indeed simultaneous to the increase in the accumulation of miR845 members during pollen development, there is a parallel decrease of 24 nt sRNAs from their targets and a progressive increase of 21/22 nt sRNAs (Figure 5d-e), potentially a consequence of their targeting.

Nevertheless, this needs to be tested since our PARE sequencing excluded non-polyadenylated RNAs. Interestingly, the preferential loading of miR845 by AGO5 correlates with low levels of CHH methylation in the SCs^51^. Due to the proposed role of miR845 in targeting of Pol IV transcripts^9^, we speculate that increased presence and activity of this miRNA in the SCs upon AGO5 loading impairs CHH methylation establishment. Interestingly, both AGO1 and AGO5 are able to weakly rescue the triploid block-induced seed collapse (Figure 3d-e), which might be the consequence of their redundant ability to load miR845 family members.

Together with this, our PARE sequencing and analysis has identified a series of transposons that were transcribed by Pol II and regulated by miRNAs (Figure 6). These transposons are likely regulated primarily by the RdDM pathway, due to their strong loss of 24 nt sRNAs and low values of CHH methylation in the unicellular stage (Figure 6c-f). Furthermore, miRNA-targeted TEs in the pollen grain seemed to not DME-mediated demethylation in the VN since they keep significantly higher CG methylation levels compared to the rest of TEs in the VN and their C-methylation values in the VN are not affected in a *DME* mutant (Figure 6f-h). We speculate that miRNA-targeting for these TEs might be a safeguard mechanism to avoid their spurious expression.

In summary, our work highlights the relevance of miRNAs for the developmental and epigenetic events that occur during the pollen grain development. The pollen grain needs to face the duality of reprogramming the transcriptome and epigenome of the newly established gametes in the sperm cells, while accomplishing a complex developmental program that culminates in the germination of the pollen tube and the successful transfer of the male gametes to the female gametophyte. Like in plants, in mouse and human cell lines changes in DNA methylation and miRNAs are an important part of the reprogramming of cells to pluripotency ^53-55^, which might be also linked to a potential miRNA control of DNA methylation, cell cycle transitions and regulation of apoptosis ^56,57^. Our work highlights that the complexity of the orchestration of the miRNome is not exclusive of mammalian reprograming for pluripotency, but also takes place during reproductive reprogramming in plants.

## Methods

### Plant material

Plants were grown under standard long day conditions at 22 °C. The mutant alleles used in this study were *ein3-1* (NASC accession number: N8052), *sks12-1* (SALK_061973) and *ago1-27*. The *KRP6pro:P19-RFP* transgene construction, plant transformation and selection were performed as described in Martinez et al (2016). Primers used for cloning are shown in Supplementary Table 6.

### Total RNA, sRNA Northern blot AGO immunoprecipitation and sRNA/PARE library construction

Total RNA was isolated using TRIzol reagent (Life Technologies). For microRNA Northern blot detection, 5 µg of total RNA were loaded in each lane for pollen developmental stage Northern blots. sRNA gel electrophoresis, blotting, and cross-linking were performed as described in Pall *et al*. (2008)^58^. The AGO1 and AGO5 proteins were immunoprecipitated using commercially available polyclonal AGO1 and AGO5 antibodies (Agrisera AB). AGO immunoprecipitated sRNA libraries were constructed as indicated in McCue et al (2012) adapted to pollen tissue. PARE libraries were constructed following the protocol described in Zhai et al (2014) adapted to pollen tissue. All sRNA libraries were made using the NEBNext Small RNA Library Prep Set for Illumina (New England Biolabs) following the manufacturer instructions and using gel-enriched sRNAs as described in Martinez *et al* (2016).

### Pollen grain separation, germination, viability test and microscopy

Pollen grain separation was performed as described in Dupl’akova et al (2016). The pollen developmental stages used for sRNA sequencing correspond to the fractions termed B1 (Unicellular), B3 (Bicellular), A3 (Tricellular). Pollen germination was determined using the media recipe from Rodriguez-Enriquez et al (2013)^59^. Each germination assay was performed in triplicates. Standard Alexander staining method was used to visualize pollen grain abortion as described in Alexander MP (1969)^60^. Visualization of pollen grain germination and Alexander stained pollen grains was performed in a Leica DM RX microscope. For pollen grain fluorescence, pollen grains of T3 plants were mounted on slides containing 50% glycerol and analyzed under a Zeiss Axioplan or a Leica DMI 4000 microscope fluorescence microscopes.

### Bioinformatic analysis

sRNA libraries were trimmed using Trim Galore. Reads were aligned using bowtie with the command “bowtie –f –t –v2” that allows two mismatches. The TAIR10 version of the *Arabidopsis* genome and the miRbase version 21 were used in this analysis. Reads were normalized to reads per million to the total reads mapped to the Arabidopsis chromosomes. For PARE library analysis, miRNA cleavage events were identified using PARESnip^61^. For genome-wide plots of PARE reads, PARE libraries were aligned using bowtie to retain only perfectly matched reads (0 mismatches). The pericentromeric region limits was determined using the description from Copenhaver et al., (1999)^62^.

## Acknowledgements

The authors thank Professor Feng Qu for the *P19* clone and Professor Michael Lenhard for his gift of the cellulosic membranes used for pollen grain germination. Sequencing was performed by the SNP&SEQ Technology Platform in Uppsala. The facility is part of the National Genomics Infrastructure (NGI) Sweden and Science for Life Laboratory. The SNP&SEQ Platform is also supported by the Swedish Research Council and the Knut and Alice Wallenberg Foundation.

## Author information

### Data access

Data can be accessed at the GEO accession number with the reviewer token:

## Supplementary information

**Supplementary Figure 1.**
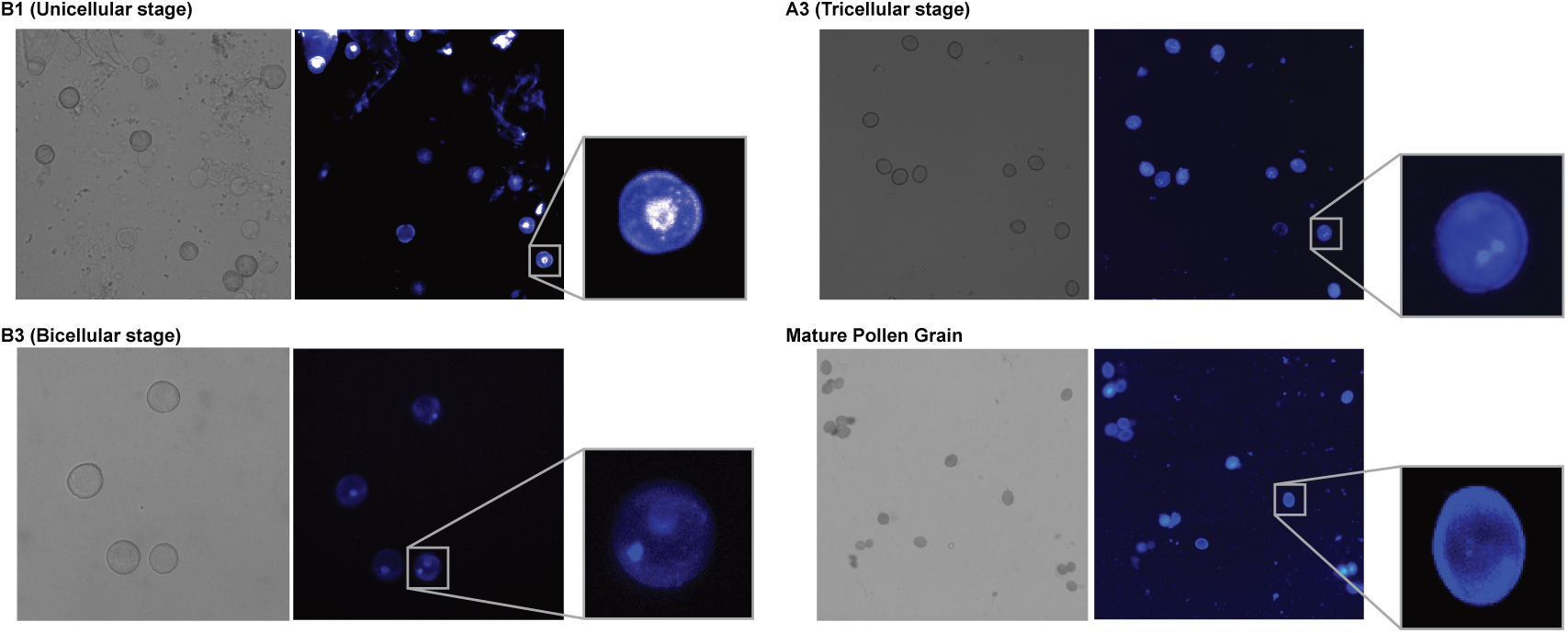
Representative pictures for each of the fractions of pollen developmental stages analyzed by sRNA high-throughput sequencing: B1 (Unicellular), B3 (Bicellular), A3 (Tricellular) and mature pollen grains.

**Supplementary Figure 2.**
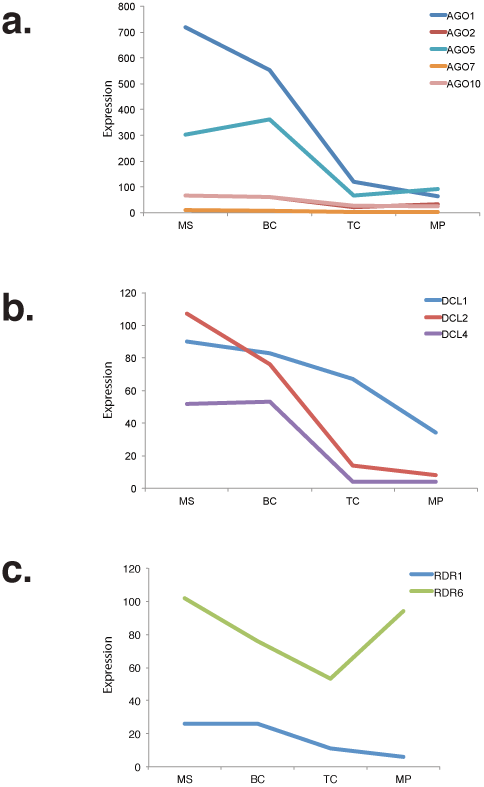
Expression pattern of different RNA silencing components involved in miRNA-related pathways during pollen development: a) AGO, b) DCL and c) RDR genes. Data extracted from ATH1 microarray. MS= microspore, BC=bicellular pollen, TC= tricellular pollen and MP= mature pollen grain.

**Supplementary Figure 3.**
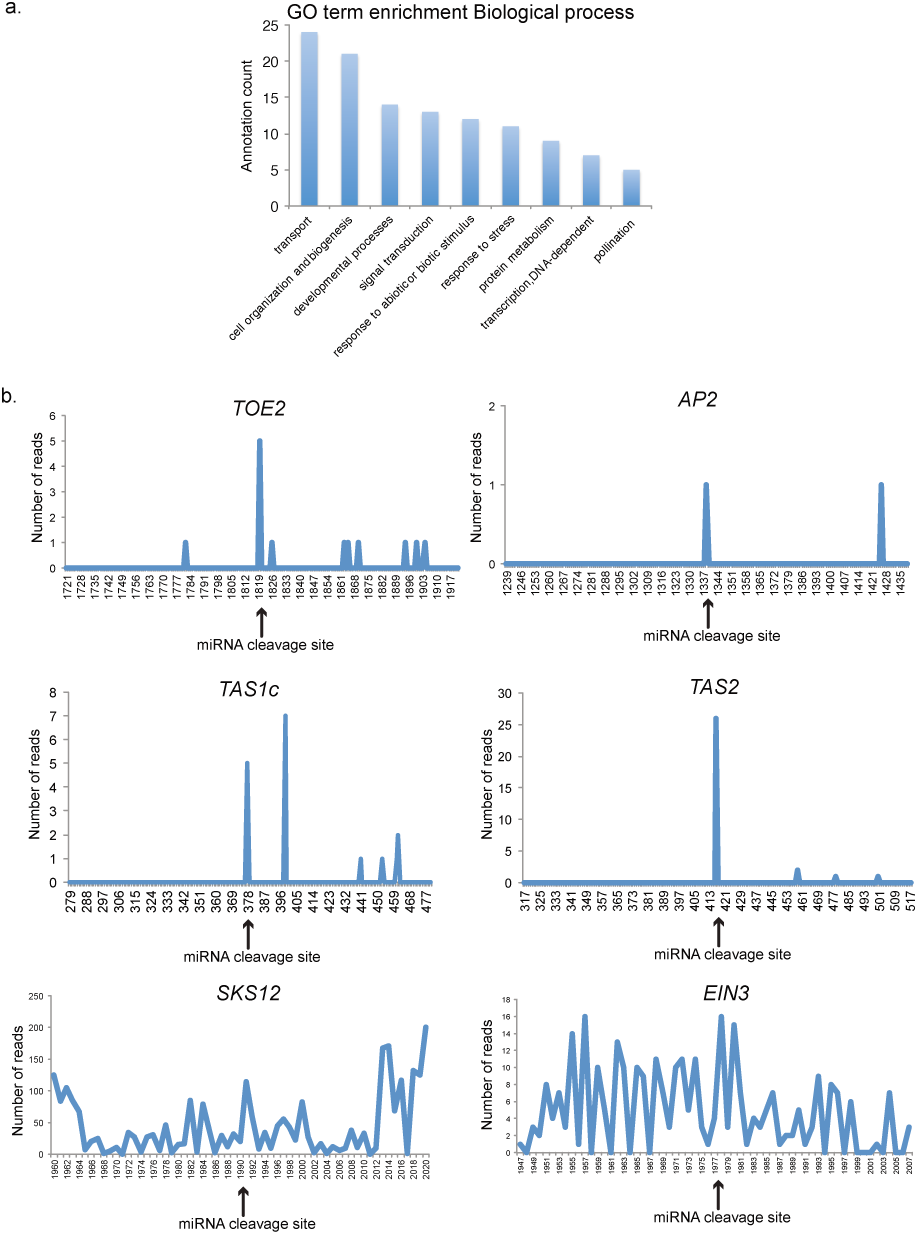
a) Analysis of the enriched GO categories for pollen specific miRNA-targeted genes. b) PARE read distribution along miRNA target sites for representative miRNA-targeted genes in Arabidopsis: miR172-targeted genes *TOE2* and *AP2* and miR173-targeted *TAS1c* and *TAS2* and the miRNA-targeted genes analyzed here: *SKS12* and *EIN3*.

**Supplementary Figure 4.**
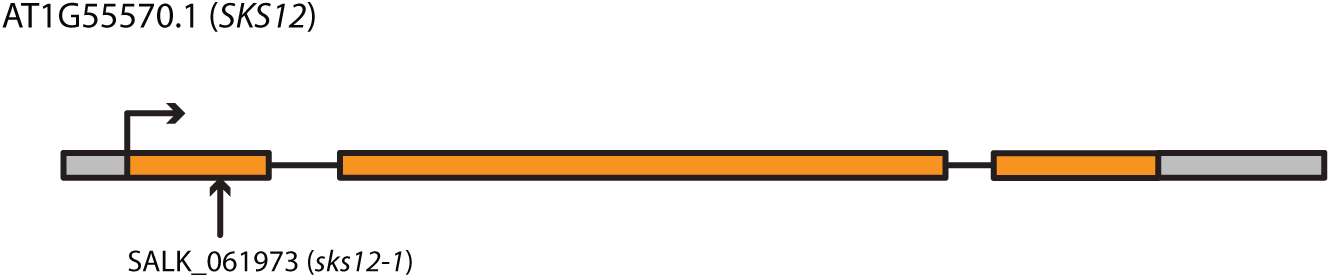
Diagram showing the location of the T-DNA insertion for the *sks12-1* mutant analyzed in this study (SALK_061973).

**Supplementary Figure 5.**
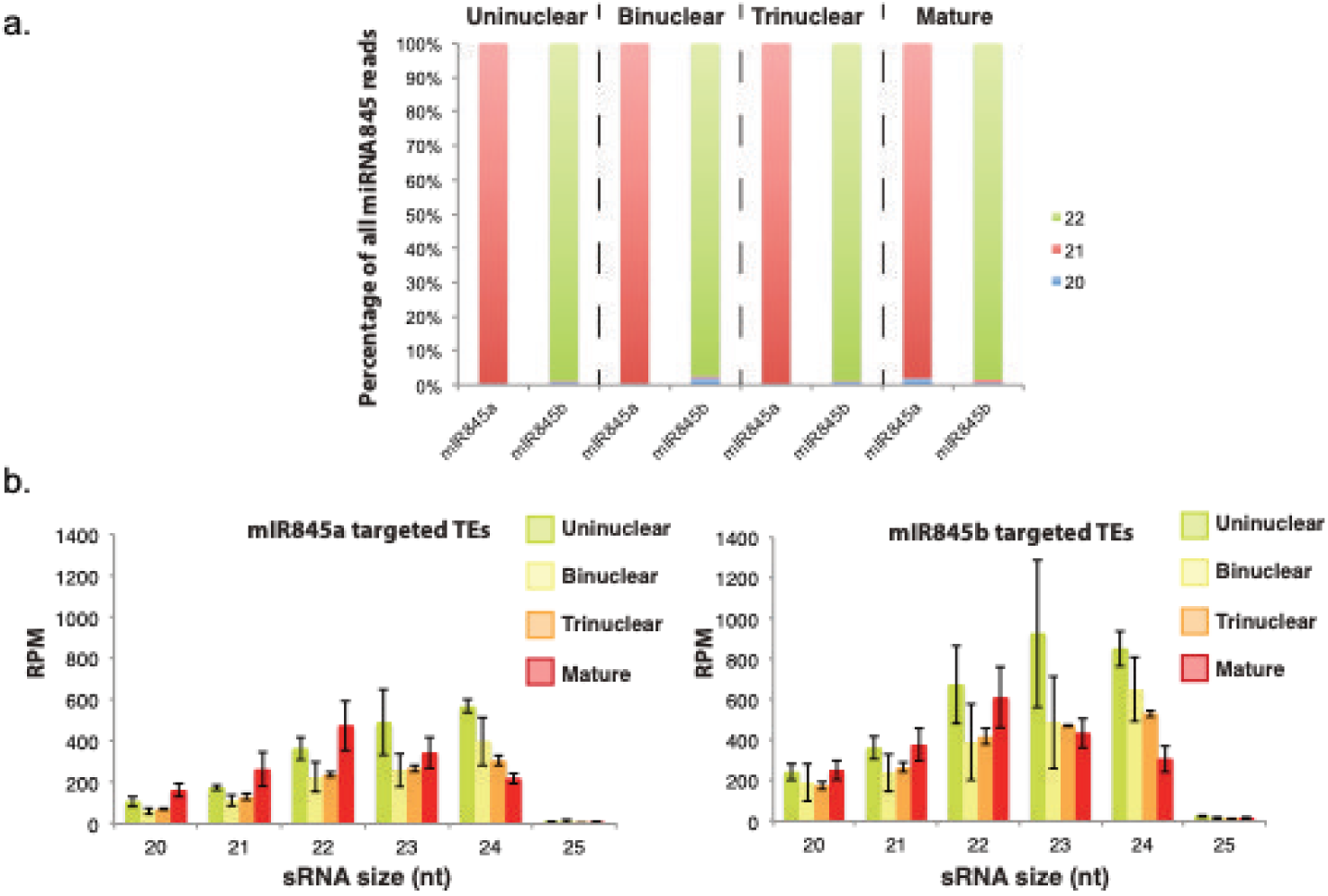
Analysis of miR845 family members and target TEs during pollen development. a) Preferential sRNA size for miR845a and b during pollen development. b) Accumulation profile of miR845a- and miR845b-targeted TEs.

**Supplementary Figure 6.**
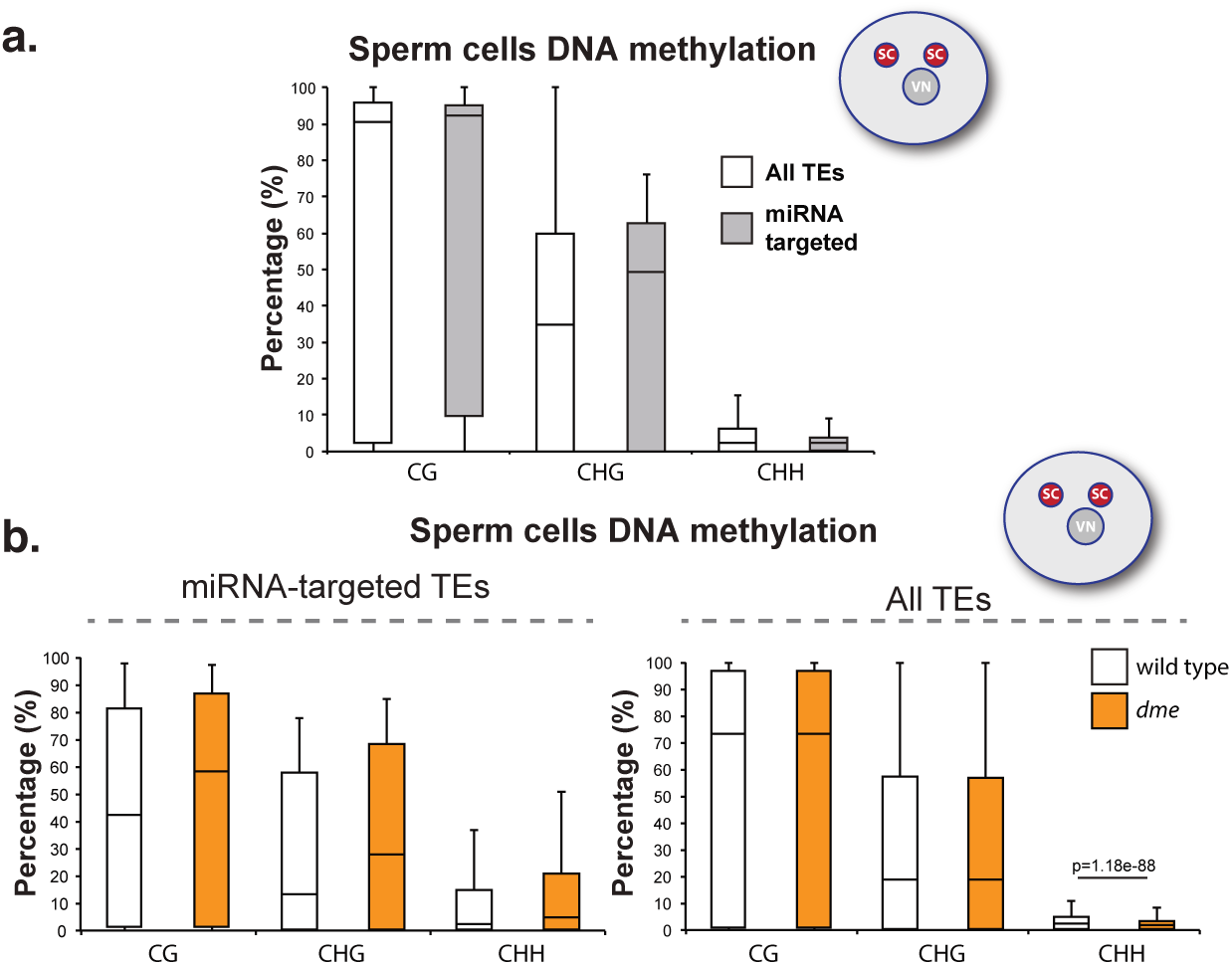
a) Levels of cytosine methylation for the different contexts (CG, CHG and CHH) in the SCs for all TEs (white boxes) or miRNA-targeted TEs (grey boxes). b) Levels of cytosine methylation for the different contexts (CG, CHG and CHH) in the SCs for miRNA-targeted TEs (left panel) or all TEs (right panel) in wild type (white boxes) and *dme* (orange boxes). Whiskers in the box plots extent to the maximum and minimum values.

**Supplementary Figure 7.**
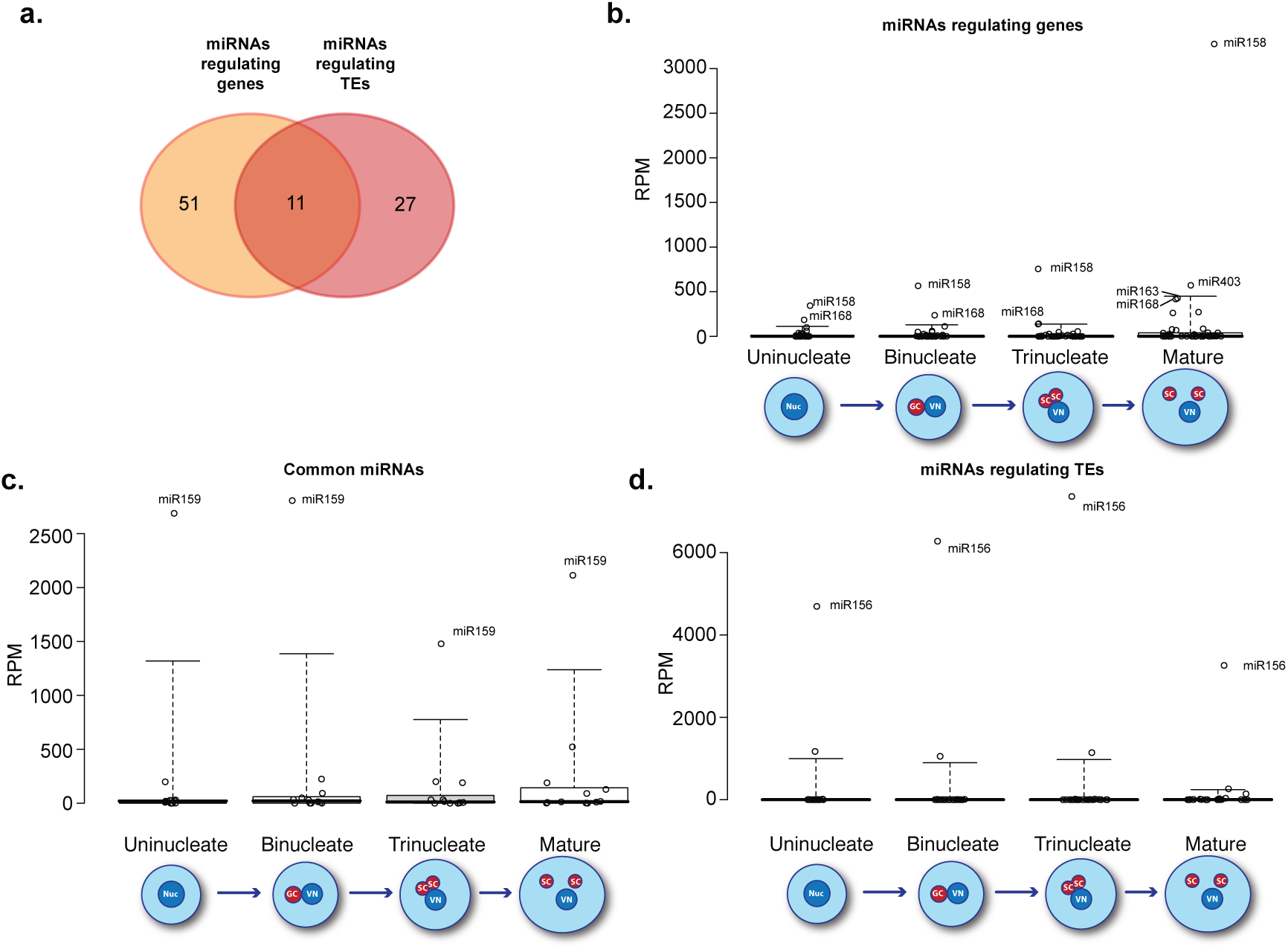
a) Venn diagram depicting the overlap between the number of PARE-identifed miRNAs regulating genes and TEs. b-d) Box plots showing the accumulation values in reads per million (RPM) of PARE-identified miRNAs regulating genes (b), TEs (d) or both (c). Whiskers in the box plots extent to the 5^th^ and 95^th^ percentile.

**Supplementary Figure 8.**
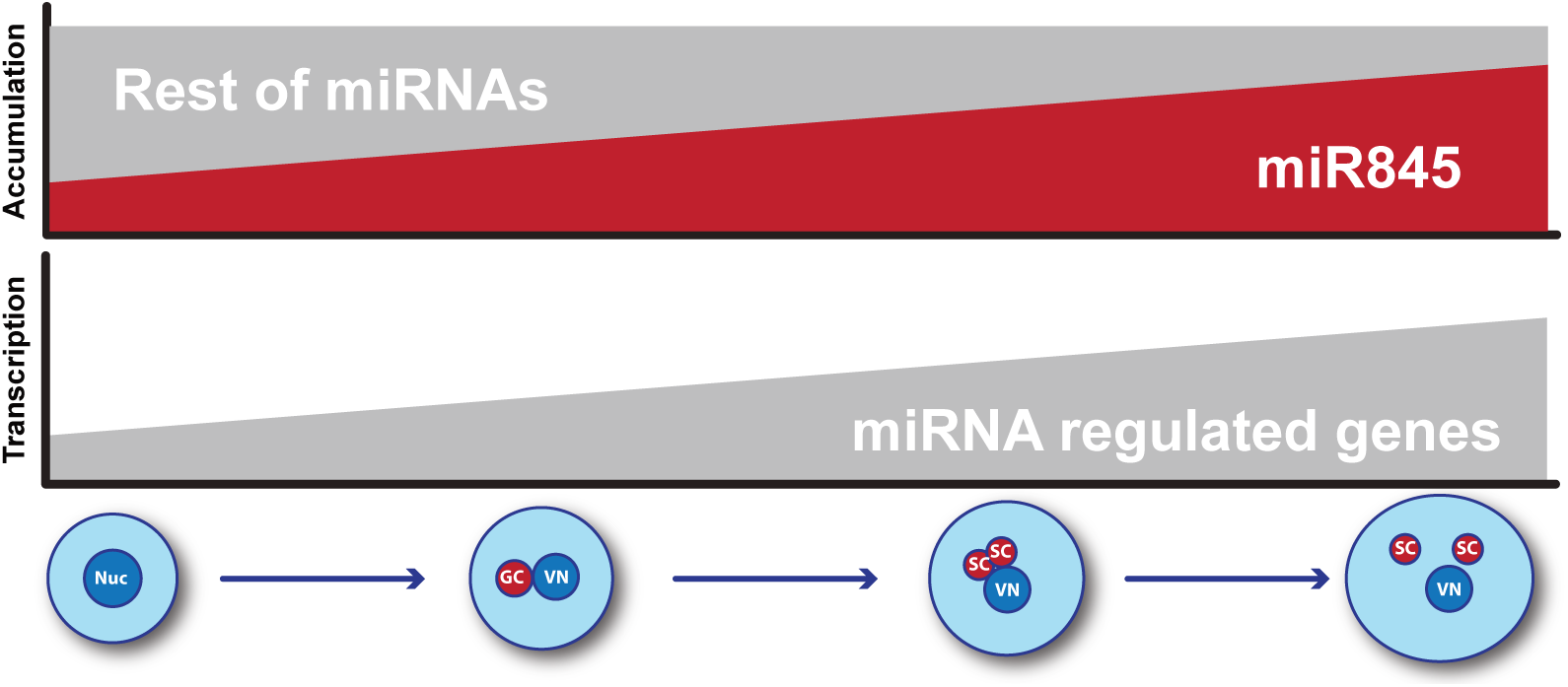
Graphic conclusion. During pollen grain development the miRNome is reprogrammed to overload miR845 members at maturity.

**Supplementary Table 1**. Libraries produced in this study.

**Supplementary Table 2**. Common and tissue specific miRNAs identified in this study.

**Supplementary Table 3**. Pollen miRNA-targeted genes identified by PARE sequencing.

**Supplementary Table 4**. Pollen miRNA-targeted TEs identified by PARE sequencing.

**Supplementary Table 5**. Publicly available data analyzed in this study.

**Supplementary Table 6**. Primers used in this study.

